# Neural contrast sensitivity is not affected by myopic blur

**DOI:** 10.1101/2024.08.30.610275

**Authors:** Niklas Domdei, Jonas Müller, Lisa Renner, Julius Ameln, Katharina Breher, Wolf Harmening, Siegfried Wahl

**Author notes:** **Corresponding author:** Siegfried Wahl. **Declaration of interests N. Domdei**, Carl Zeiss Vision International GmbH (E); **J. Müller**, None; **L. Renner**, None; **J. Ameln**, None; **K. Breher**, Carl Zeiss Vision International GmbH (E); **W.M. Harmening**, None; **S. Wahl**, Carl Zeiss Vision International GmbH (E). The funders did not have any additional role in the study design, data collection and analysis, decision to publish or preparation of the manuscript.

## Abstract

**Purpose:** The prevalence of myopia is increasing worldwide, accompanied by an increase of potentially under-corrected myopes. Because the neural pathways processing the retinal image are prone to adaptation in relation to the retinal image quality, we wondered to what extent neural contrast sensitivity (NCS) is altered in the presence of myopic blur. Additionally, the impact of retinal abnormalities like foveal hypoplasia with albinism on NCS was tested.

**Methods:** NCS was psychophysically determined for 11 emmetropic, 23 myopic well-corrected and 15 myopic under-corrected otherwise healthy young (27 ± 6 years) participants and 1 albinism patient. Aberration-free stimulation, independent of the eye’s refractive state, was achieved by using a unique spatial light modulator-based interferometric system to bypass the eye’s optics.

**Results:** No significant differences in NCS were observed between the three groups (Median area-under-curve: 61.9, 62.1, and 62.9 for emmetropes, well-corrected, and under-corrected myopes, respectively; all p > 0.1) but were significantly equivalent between emmetropes and myopes (all p < 0.001). However, the NCS function of the albinism patient differed significantly from the here defined “normal” NCS function.

**Conclusions:** NCS is unaffected by myopic blur and remains stable even for under-correction of up to 1.5 D. This means, that long-term under-corrected myopes still can achieve normal visual acuity as soon as their refractive errors are sufficiently corrected. Furthermore, NCS testing can relate visual deficits to an underlying neurological disorder.

## Introduction

Myopia, a common eye condition in humans with increasing prevalence worldwide,^1,2^ leads to a blurred retinal image if inadequately corrected. As known from amblyopia, uncorrected refractive errors can result in slight to severe visual impairment if left untreated.^3^ According to a recent prospective, almost 10 % of the Australian population is under-corrected and could achieve an increase in visual acuity of at least one line on the visual chart with adequate correction.^4^ Due to the increasing cases of myopia, it can be assumed that the number of under-corrected eyes has also grown and will be growing even further in the future. Additionally, purposefully induced under-correction is also discussed as a possible myopia progression strategy.^5^

Spatial vision, and specifically, our ability to detect visual contrast, is a product of optical and neural processes that start with a retinal image in the eye and ultimately lead to visual perception. The optical properties of the eye that determine the image quality formed on the retina,^6,7^ can be mathematically described by the modulation transfer function (MTF). On the neural end, the neural transfer function (NTF) describes the processing of that image by the retina and ascending visual pathways.^8^ The psychophysically measurable ability to detect contrast, the contrast sensitivity function (CSF), can thus be written as,

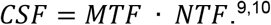

Direct measurement of the CSF is achieved by testing the minimum visible contrast at varying spatial frequencies.^11^ However, if a patient’s CSF deviates from the basic healthy CSF, it is often unclear whether this is the result of impaired optics (MTF) or neural pathways (NTF). Furthermore, it was observed that a deterioration of the eye’s optical components is followed by a degradation of the neural components, according to the use-it-or-lose-it principle.^12^

For example, by using adaptive optics to eliminate any MTF related influences,^13–15^ one specific component of the NTF, the neural contrast sensitivity (NCS), could be measured directly in patients suffering from keratoconus, characterized by an extremely blurry retinal image due to highly irregular corneal aberrations. Keratoconus patients were unable to achieve normal NCS.^16^ It was argued that due to the unusually high optical aberrations, which attenuate high spatial frequencies in particular, the missing input power of such frequencies lead to a degradation of the processing channels sensitive to high spatial frequencies.

Direct measurement of the NCS can be achieved with an interferometric approach that focusses two coherent light beams into the eye’s pupil, producing the interference fringes stimulus on the retina.^17–19^ In this so called Maxwellian view configuration,^20^ the eye’s optics are bypassed and an aberration-free presentation of the stimulus is achieved. Such a system could furthermore be beneficial to assess the underlying reason of poor visual performance. For example, if significantly reduced NCS values are observed for a patient, it can be assumed that vision is limited due to a neurological or retinal disorder instead of optical properties.

Here, we tested whether the blur that is present in myopic eyes, and in particular in cases of chronically under-corrected myopia, leads to a reduced NCS. We used a spatial-light-modulator-based interferometer instrument to study the NCS of emmetropic, well-corrected and under-corrected myopes. Additionally, the resulting eye-healthy NCS is compared with the NCS of an albinism patient with known congenital foveal hypoplasia, thereby probing the method’s capability to detect an estimated abnormal NCS.

## Methods

The study adhered to the tenets of the Declaration of Helsinki. The ethics authorization to perform the measurements was granted by the Medicine Faculty Human Research Ethics Committee from the University of Tübingen (616/2022BO2 and 094/2024BO1). Before data collection, the experiment was explained in detail to the participants, and written informed consent was collected from each participant. All data were pseudonymized and stored in full compliance with the principles of the Data Protection Act GDPR 2016/679 of the European Union.

### Study participants and group assignment

A total of 49 healthy participants (28 females; 27 ± 6 years) were recruited, all students at the University of Tübingen with no known retinal pathologies. Additionally, one otherwise healthy Albinism patient (f; 31 years) with known foveal hypoplasia was recruited.

At the first visit, each participant underwent objective refraction (iProfiler plus, Carl Zeiss Vision GmbH, Aalen, Germany) and visual acuity testing with their current glasses, if available. Participants were assigned to the emmetropic study group if the respective spherical equivalent (SE) of the objective refraction was ≥ -0.5 D and ≤ +0.5 D and they achieved a visual acuity of -0.2 logMAR (VisuScreen 500, Carl Zeiss Vision GmbH, Aalen, Germany). Myopic participants (SE < -0.5 D) additionally underwent subjective refraction (ZEISS Visuphor 500, Carl Zeiss Vision GmbH, Aalen, Germany) to determine if the participant belonged to the well-corrected or under-corrected group. Participants were assigned to the under-corrected group if subjective refraction revealed a deviation of at least 0.5 D (SE) between worn and needed refractive aid accompanied by an increase of visual acuity by at least one line when optimally corrected.

Axial length of the tested eye was measured (IOLMaster 700, Carl Zeiss Meditec, Dublin, CA, USA), and to ensure retinal integrity, foveolar optical coherence tomography (OCT; ZEISS PlexElite 9000, Carl Zeiss Meditec, Dublin, CA, USA) scans were recorded for each participant.

### Interferometer setup

To investigate the neural contrast sensitivity of the different groups, the threshold of contrast vision was measured psychophysically using a liquid-crystal-on-silicon spatial-light-modulator-based interferometer (Fig. 1A) as described earlier.^21^ In brief, the necessary spatially coherent light in this setup was provided by a supercontinuum laser (NKT compact, SuperK Compact, NKT Photonics, Birkerød, Denmark) and the used narrowband green light (550 ± 5 nm; FBH550-10, Thorlabs GmbH, Bergkirchen, Germany) was filtered from its broad spectrum. The key element of the system, the spatial light modulator (PLUTO-2-VIS-016, Holoeye, Berlin, Germany), was placed in the Fourier plane of the first collimating lens. To create the two laterally separated coherent wavefronts the spatial light modulator displayed two blazed gratings, with each grating providing a tilt that controlled the lateral shift of the wavefront in the image plane of the system. A spherical mirror focused the two beams in the system’s intermediate image plane, where a motorized iris diaphragm (8MID10-40, Standa Ltd., Vilnius, Lithuania) was placed as a field stop to filter out zeroth and higher diffraction orders. Additionally, a tunable lens (EL-12-30-TC, Optotune Switzerland AG, Dietikon, Switzerland) was placed at this position to compensate for the doubling of the field stop in cases of uncorrected refractive error resulting in non-overlapping beams on the retina^22^. The power of the tunable lens was set individually by each participant at the beginning of the experiment to completely overlap the two spots on the retina. After collimating the wavefront again, the wavefront was focused together with a non-coherent background beam in the pupil plane. The resulting Maxwellian field of view was 1.5° for the stimulus and 1.6° for the background. The non-coherent background was necessary to decrease the available contrast values into the range of the expected thresholds and to reduce masking effects of coherent spatial noise due to the monochromatic light. Spatial coherence of the background was broken via a rotating diffuser foil. The incoherent background light accounted for 90% of the total retinal illuminance,^23,24^ which was about 300 Td for the combined field.^23,25^

**Figure 1.**
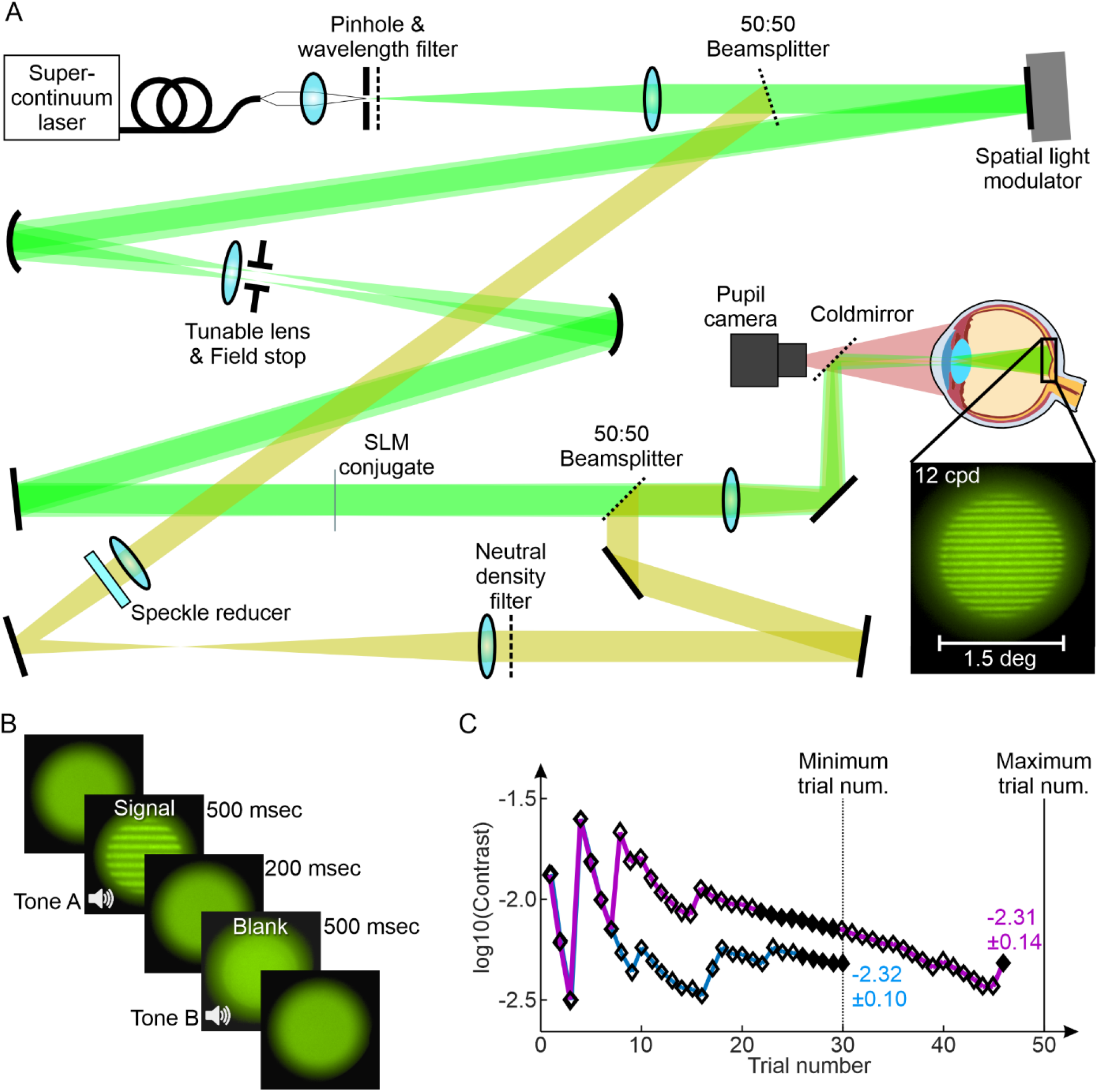
The interferometric setup and psychophysical procedure for the neural contrast sensitivity assessment. A) Schematic beam path of the optical system forming the interference fringes on the retina by using a spatial light modulator (SLM). A motorized aperture adjusts the diameter of the field stop blocking higher order maxima as a result of the SLM based approach splitting the light source into two via a phase mask. The background consisted of the same light spectrum as the stimulus (indicated here with a yellowish path) and was used for a better contrast control as the background was always visible with the stimulus being projected on top of the background light. The tunable lens was adjusted by each participant to have the two spots perfectly overlapping on the retina. An on-axis pupil camera was used to ensure optimal stimulus delivery throughout the experiment. B) Each trial consisted of two intervals and was started by the participant. One interval contained the stimulus, the other one a blank and the participant had to report the stimulus (signal) interval via a button press (2-Interval forced choice). The two intervals were accompanied by two tones and separated by an inter stimulus interval of 200 msec. C) Exemplary experiment progression with QUEST for two runs. Stop criteria were a confidence interval for the current threshold estimate smaller than 0.15 (indicated by filled markers) and a minimum number of 30 trials. If the first criterion was not met after the first 30 trials, additional trials (up to 50 in total) were presented until a SD <0.15 was reached. Runs not meeting the criterion after 50 trials were repeated.

Because even small head movements would move the pupil out of the beam, the participants were asked to bite on an individually produced bite bar during the measurements. The eye’s pupil was then conjugated with the system’s pupil position via a x-y-z-translation stage, monitored in real-time throughout the experiment by an on-axis CMOS camera (DMK 27AUP031, The Imaging Source, Bremen, Germany) behind a cold mirror (FM203, Thorlabs GmbH, Bergkirchen, Germany). Testing only dominant eyes, the fellow eye was covered with an eye patch during the NCS measurement.

### Psychophysical measurement of the neural contrast sensitivity

The NCS measurement procedure was conducted via QUEST (Quick estimate by sequential testing),^26^ an adaptive staircase method for threshold estimation, in a two-interval forced choice environment (Fig. 1B) using MATLAB (The MathWorks, Inc., Natick, USA) and the psychophysics toolbox.^27^ The participant started each trial individually. After a delay of 300 msec, the first of the two intervals was presented for 500 msec, followed by an interstimulus interval of 200 msec and the second interval (again 500 msec). With the background always visible, the two intervals were projected on top of it accompanied by audio cues helping the participant to separate the two intervals. Signal and blank were assigned randomly into the two intervals. The participant named the interval in which the grating had been perceived via a button press on a controller. Depending on the answer being correct or incorrect the contrast of the next stimulus changed as specified by QUEST. Every 5^th^ trial was either a “catch” trial, displaying a blank in both intervals or a “lapse” trial with maximum contrast in turn. For each run, the participant had to do at least 30 trials. If QUEST ’s confidence interval for the current threshold estimate was smaller than 0.15, the run was finished, otherwise more trials, up to a maximum number of 50 trials, had to be done until the before mentioned criterion was met (see Figure 1C). If the stop criterion was not met after 50 trials the run was terminated and had to be repeated. On average runs were completed with 31.78 ± 3.64 trials, with 95% of trials being completed after 39 trials.

Participants had to do two training runs at the beginning of the experiment with 12 and 24 cpd to familiarize themselves with the unusual stimulus presentation and experiment procedure. In general, a total of seven different spatial frequencies (SF) were measured across three runs, which include 3, 6, 12, 18, 24, 30 and 36 cpd. A subgroup of emmetropes (N = 8) was tested additionally at 9 cpd for a more detailed sampling of the NCSF. SFs were tested in pseudo-randomized order to minimize bias from training effects or fatigue. After completion of a set of SFs, a short break (about 5 min) was taken. Each SF was tested three times. If the difference between the lowest and highest threshold was greater than 0.2, a fourth and, if needed, a fifth run for this SF was recorded. This limit of 0.2 was determined empirically during pilot testing based on the observation that repeated measurements at the same SF fluctuate with about ±0.1. Fluctuation of 0.2 and higher could only be observed during training runs or the participant being inattentive. The final here reported NCS at a given SF for each participant, was calculated as the median of the three closest recordings.

### Adaptive optics retinal imaging and foveolar cone mosaic analysis

The central ± 150 µm in the dominant eyes of three selected participants were imaged using near-infrared light for imaging and wavefront sensing, filtered dichroically (788 ± 12 nm; FF01-788/12-25, Semrock, Rochester, NY, USA) from the output of a supercontinuum laser light source (SuperK EXTREME, NKT Photonics, Birkerød, Denmark). Adaptive optics correction, run in a closed loop at about 25 Hz, consisted of a Shack–Hartmann wavefront sensor (SHSCam AR-S-150-GE; Optocraft GmbH, Erlangen, Germany) and a 97-actuator deformable mirror (DM97-08; ALPAO, Montbonnot-Saint-Martin, France) placed at a pupil conjugate. The imaging raster spanned a square field of 0.85 × 0.85 degrees of visual angle. The light reflected from the retina was detected in a photomultiplier tube (PMT, H4711-50, Hamamatsu, Japan), located behind a confocal pinhole (0.5 Airy disk diameter). PMT signals were sampled by a field programmable gate array (FPGA) board (ML506; Xilinx, San Jose, CA, USA), producing video frames with 512 × 512 pixels (spatial resolution, 0.1 arcmin of visual angle per pixel) at about 27 or 30 Hz. To ensure optimal image quality during recording, the pupil’s position relative to the adaptive optics scanning laser ophthalmoscope (AOSLO) beam was carefully maintained.^28^ Videos were recorded at the preferred retinal locus of fixation. Optimal image quality was found by selecting the best video from five to ten PRL-centered videos recorded using different defocus settings of the deformable mirror. All videos were 10 seconds long. Acquired AOSLO video frames were spatially stabilized by offline, strip-wise image registration using a modified version of previously published software in Matlab.

The processing pipeline to determine the cone density centroid was described earlier^29^. In brief, the cone center locations in the final montage were labeled in a semi-manual process by a single trained image grader: first, a convolutional neural network was used to annotate retinal images automatically^30,31^ and in a second step manually corrected using custom software in Matlab. Based on the labeled cone center locations, a Voronoi tessellation was computed (Matlab: delaunayTriangulation, voronoiDiagram and voronoin). Each cone was regarded as occupying the space of each corresponding Voronoi cell. Angular cone density (cones/deg^2^) was computed at each image pixel by averaging the combined Voronoi area of the nearest 150 encircled cones around that pixel. Finally, the cone density centroid (CDC) was determined as the weighted centroid (Matlab: regionprops (‘WeightedCentroid’)) of the highest 20% of cone density values. The participant’s Nyquist limit in term of cut-off frequency was calculated as 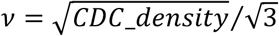.^32^

### Visual acuity testing and estimation

For a more direct assessment of the best corrected minimum angle of resolution, two selected participants and the albinism patient underwent visual acuity testing with the Freiburg visual acuity test (FrACT) using the Landolt C in 8 orientations.^33^ The screen was placed at a 3.5 m distance and calibrated according to the instructions. The final value reported here for the FrACT visual acuity was the average of three consecutive measurements.

To estimate the albinism patient’s visual acuity based on retinal anatomy, the approach and equation proposed by Woerz et al., 2024 were utilized.^34^ To that end, spectral-domain OCT images of the fovea were recorded using a prototype high-resolution device (High-Res OCT, Heidelberg Engineering GmbH, Heidelberg, Germany) with an axial resolution of about 2 µm (in air). The required band ratios were evaluated in the foveolar B-Scan defined by the B-Scan showing the deepest foveal indentation.

### Quantification and statistical analysis

All numerical and statistical analysis was performed in Matlab. The area under the curve was calculated as the numerical integration (Matlab: trapz). Normal distribution was tested via Kolmogorov-Smirnov test (Matlab: kstest). Because data was not normally distributed, the three groups were compared statistically with Wilcoxon’s rank sum test (Matlab: ranksum), after confirmation of homoscedasticity via Brown–Forsythe test for equality of variances (Matlab: vartestn(‘BrownForsythe’)). Additionally, to specifically test for similarity of the found NCS functions, an equivalence test based on the two one-sided tests approach (TOST) was implemented using the Mann-Whitney U-test test.^35,36^ Correlations between NCS and the eye’s refractive state were calculated based on the F-test (Matlab: regress) and confidence intervals computed using the functions: fitlm and predict. The violin plots were created with Holger Hoffmann’s Matlab function.^37^

## Results

Neural contrast sensitivity (NCS) was psychophysically assessed in the dominant eyes of a total of 49 participants by projection of interference fringes, a method which is known not to be affected by individual optical errors. Participants were assigned into three different groups based on the participant’s currently needed and available optical correction (Table 1). The first group consisted of emmetropes (control) defined by an objectively measured spherical equivalent of ≤ ±0.5 D in spherical equivalent (N = 11). The second group were the well-corrected myopes (N = 23) defined by a maximum difference of ±0.5 D (spherical equivalent) between needed and available correction. The third group, the under-corrected myopes (N = 15), was defined by an offset of > 0.5 D (sphere or cylinder) between needed and available correction and an improvement in visual acuity by at least one line on the visual acuity chart when best-corrected.

**Table 1.** Demographics, refraction and visual acuity of all participants sorted by study groups. In addition to the best-corrected visual acuity (BCVA), visual acuity (right column) was tested for myopes with their current spectacle correction.

| ID | Sex | Age | Axial length (mm) | Objective refraction (SE in Diopter) | BCVA (LogMAR) | Spectacle correction (SE in diopter) | Acuity with spectacles (LogMAR) |
| --- | --- | --- | --- | --- | --- | --- | --- |
| Emmetrop_001 | m | 24 | 23.9 | -0.42 | -0.2 | - | - |
| Emmetrop_002 | f | 23 | 22.7 | -0.45 | -0.2 | - | - |
| Emmetrop_003 | f | 25 | 23.4 | -0.49 | -0.2 | - | - |
| Emmetrop_004 | f | 24 | 24.0 | -0.18 | -0.2 | - | - |
| Emmetrop_005 | f | 29 | 22.7 | 0.36 | -0.2 | - | - |
| Emmetrop_006 | m | 30 | 22.6 | -0.27 | -0.2 | - | - |
| Emmetrop_007 | f | 33 | 23.7 | -0.51 | -0.2 | - | - |
| Emmetrop_008 | f | 21 | 23.6 | -0.38 | -0.2 | - | - |
| Emmetrop_009 | f | 28 | 25.3 | 0.38 | -0.2 | - | - |
| Emmetrop_010 | f | 34 | 23.9 | -0.16 | -0.2 | - | - |
| Emmetrop_011 | m | 39 | 22.1 | 0.39 | -0.2 | - | - |
| MyoWell_001 | f | 29 | 24.7 | -4.45 | -0.2 | -4.750 | -0.2 |
| MyoWell_002 | f | 23 | 23.1 | -1.98 | -0.2 | -2.125 | -0.2 |
| MyoWell_003 | f | 31 | 24.1 | -2.19 | -0.2 | -1.750 | -0.2 |
| MyoWell_004 | f | 32 | 23.2 | -2.09 | -0.2 | -2.000 | -0.2 |
| MyoWell_005 | f | 22 | 24.0 | -1.98 | -0.1 | -1.125 | -0.1 |
| MyoWell_006 | f | 25 | 23.7 | -1.76 | -0.2 | -1.625 | -0.2 |
| MyoWell_007 | m | 27 | 23.7 | -2.13 | -0.2 | -1.500 | -0.2 |
| MyoWell_008 | m | 28 | 26.5 | -4.08 | -0.2 | -4.000 | -0.2 |
| MyoWell_009 | f | 35 | 23.9 | -2.15 | -0.2 | -1.750 | -0.2 |
| MyoWell_010 | f | 23 | 24.2 | -0.85 | -0.1 | -1.000 | -0.1 |
| MyoWell_011 | m | 37 | 25.1 | -2.01 | -0.2 | -2.000 | -0.2 |
| MyoWell_012 | m | 22 | 23.7 | -1.94 | -0.2 | -1.875 | -0.2 |
| MyoWell_013 | f | 20 | 22.7 | -1.36 | -0.1 | -1.500 | -0.1 |
| MyoWell_014 | f | 23 | 24.2 | -1.94 | -0.1 | -1.500 | -0.1 |
| MyoWell_015 | m | 22 | 23.2 | -1.56 | -0.2 | -1.750 | -0.2 |
| MyoWell_016 | f | 39 | 24.6 | -2.72 | -0.2 | -3.125 | -0.2 |
| MyoWell_017 | m | 24 | 24.0 | -4.92 | -0.2 | -4.250 | -0.2 |
| MyoWell_018 | m | 30 | 24.0 | -2.44 | -0.2 | -2.625 | -0.2 |
| MyoWell_019 | m | 29 | 26.5 | -4.14 | -0.2 | -3.750 | -0.2 |
| MyoWell_020 | f | 29 | 24.7 | -3.40 | -0.2 | -2.875 | -0.2 |
| MyoWell_021 | f | 25 | 24.3 | -2.71 | -0.2 | -1.750 | -0.2 |
| MyoWell_022 | m | 36 | 24.9 | -2.13 | -0.2 | -1.875 | -0.2 |
| MyoWell_023 | f | 34 | 24.4 | -6.65 | -0.2 | -6.375 | -0.2 |
| MyoUnder_001 | m | 31 | 23.9 | -0.83 | -0.2 | none | 0 |
| MyoUnder_002 | m | 33 | 23.5 | -0.54 | -0.1 | none | 0.1 |
| MyoUnder_003 | f | 20 | 23.8 | -1.02 | -0.2 | none | 0.1 |
| MyoUnder_004 | m | 33 | 24.1 | -2.30 | -0.2 | -2.250 | 0 |
| MyoUnder_005 | f | 20 | 23.5 | -0.80 | -0.2 | -0.375 | 0 |
| MyoUnder_006 | f | 35 | 23.8 | -1.10 | -0.2 | none | 0.1 |
| MyoUnder_007 | f | 19 | 25.2 | -1.96 | -0.2 | -1.125 | 0 |
| MyoUnder_008 | f | 23 | 23.3 | -2.95 | -0.1 | -2.250 | 0 |
| **MyoUnder_009** | f | 24 | 22.9 | -0.58 | -0.2 | none | 0 |
| **MyoUnder_010** | m | 23 | 23.7 | -1.49 | -0.2 | none | 0.1 |
| **MyoUnder_011** | m | 25 | 22.6 | 0.09 | -0.2 | none | 0 |
| **MyoUnder_012** | m | 22 | 26.2 | -3.00 | -0.2 | -1.500 | 0.7 |
| **MyoUnder_013** | m | 20 | 24.4 | -0.50 | -0.2 | none | 0.1 |
| **MyoUnder_014** | m | 20 | 24.1 | -2.14 | -0.2 | -2.125 | -0.1 |
| **MyoUnder_015** | m | 27 | 26.9 | -7.49 | -0.2 | -6.875 | -0.1 |
| **Albinism_001** | f | 31 | 21.3 | 0.94 | 0.2 | 1.375 | 0.2 |

### Emmetropes and myopes have equal neural contrast sensitivity

Median NCS values -in the following reported as log10 units -were similar across all three groups (see Fig. 2A). Sensitivity increased from a spatial frequency (SF) of 3 cycles per degree (cpd) (2.15, 2.17, and 2.15; for emmetropes, well-corrected, and under-corrected myopes, respectively), over 6 cpd (2.33, 2.35, and 2.36), to 12 cpd (2.38, 2.39, and 2.39). Because peak NCS was reported earlier at about 10 cpd, a subset of emmetropic participants (N = 8) was tested at 9 cpd. For this subset, median NCS measurements were equal with their respective median NCS at 12 cpd (both 2.36). The group of under-corrected myopes reached the maximum median NCS at 18 cpd (2.34, 2.38, and 2.41). Beyond this SF, NCS slowly decreased for all three groups, measured at 24 cpd (2.21, 2.29, and 2.30), 30 cpd (2.10, 2.10, and 2.13), and 36 cpd (1.96, 1.89, and 1.83). Overall variability within the three groups was very low for SFs ≤ 24 cpd, given average interquartile ranges for individual NCS values of 0.10, 0.08, and 0.11. The smallest interquartile range was observed for NCS measurements at 12 cpd with 0.05, 0.07, and 0.05, and the highest at 36 cpd with 0.58, 0.55, and 0.70 (for emmetropes, well-corrected, and under-corrected myopes, respectively). Several participants were not able to perceive the 36 cpd interference fringes even at the highest contrast setting (N = 5 (45 %), 1 (4 %), and 2 (13 %)) making a threshold determination impossible. Based on the observation of a similar NCS values, the measurements were pooled across all participants (N = 49) to calculate the normal NCS curve via a Rational-2-1 fit, describing the NCS of young (20-39 years old) and healthy humans (Fig. 2B).

**Figure 2.**
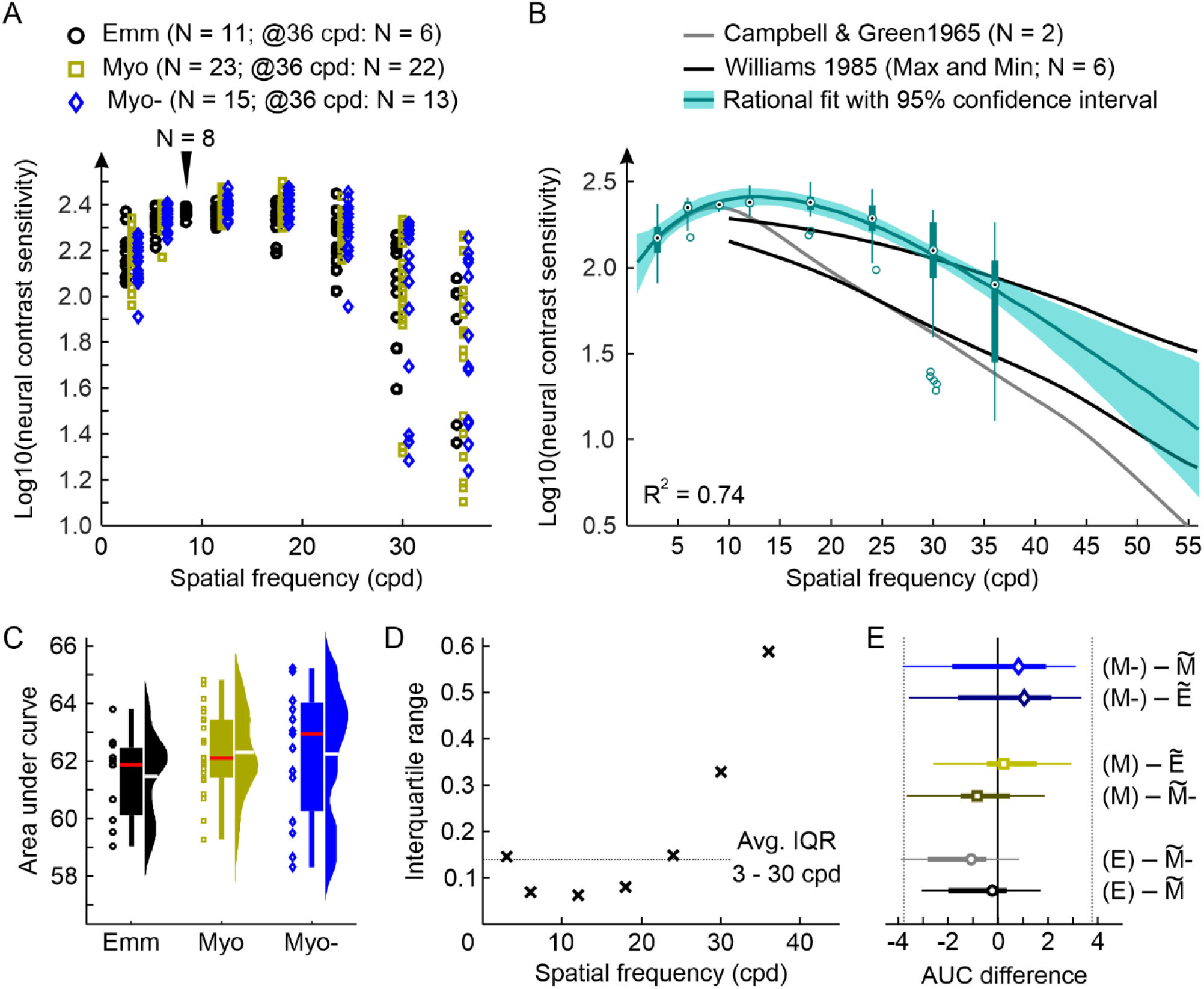
Neural contrast sensitivity measured with interference fringes and analysis. A) Log10(NCS) for the three different groups tested and comparison with literature. Marker position states the group median and error bars the interquartile range. Not all participants were able to perceive the 36-cpd-stimulus. The resulting number of valid measurements for this spatial frequency is stated in the figure legend. B) Based on the observation that NCS is similar across the different groups (see C-E for the analysis) data was pooled (plotted as compact boxplot with circles marking outliers) and fitted with a rational function (ax^2^+bx+c)/(x+d). For comparison, literature data from Campbell&Green^17^, and Williams^23^is provided. C) Area under the NCS curve from 3 to 30 cpd. White lines indicate the average, red lines the median value. Differences between groups are not significant (Mann-Whitney U-test, all p > 0.05). D) Determination of the average interquartile range per test SF from pooled thresholds for 3 to 30 cpd. E) Test for equivalence using the average IQR per tested SF as lower and upper limit (l1 = -3.01; l2 = 3.01) reveals a significant equivalence between AUCs of the three groups (TOST, all p ≤ 0.01). The thick horizontal lines display the groups interquartile range (25% -75%) with the thin lines representing the whiskers. The group’s median is given by the marker. Emmetrope: Emm, E; Myope (well): Myo, M; Myope (under): Myo-, M-; 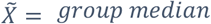.

For a statistical comparison of the recorded NCS functions, the area under the curve (AUC) was calculated for SFs between 3 and 30 cpd, neglecting 36 cpd values due to the very high variability and incompleteness of the data available (Figure 2C). The median AUC value for the emmetrope group was 61.87 [Interquartile Range: 60.30 to 62.35]. In the two myopic groups, the median AUC was 62.11 [61.48 to 63.43] for the well-corrected and 62.94 [60.66 to 63.95] for the under-corrected participants. The observed small differences between the three groups were not significant (all p > 0.1; Mann-Whitney U-test). In addition, similarity of the three groups was statistically assessed by equivalence testing following the two one-sided tests approach (TOST).^35,36^ For the TOST, it is necessary to define the limits within which any observed fluctuations are tolerable. Here, the required limit was determined based on the average interquartile range of 0.14 of NCS measurements across all participants at spatial frequencies between 3 and 30 cpd (Fig. 2D). Given this average standard deviation for NCS measurements, the AUC would change by ±3.77, which was then used as the test boundaries with L1 = -3.77 as the minimum and L2 = 3.77 maximum limit. With this limit definition, the three groups showed significantly equivalent populations of AUC for the measured NCS curves (all p ≤ 0.001, see Fig. 2E).

### Neural contrast sensitivity is independent of refractive error and age

A noteworthy observation was the increasing variability for NCS measurements at high SF (≥ 30 cpd), which was similarly high for all three groups at 36 cpd (Fig. 2A). To find an explanation for this, the individual’s median NCS at 36 cpd of all groups was tested for correlation with axial length (Fig. 3A), without revealing any significance (p = 0.48). Additionally, using the myopic eyes’ data only, the spherical equivalent and the amount of under-correction were compared with the respective individuals’ NCS at 36 cpd. Again, both correlations were not significant (both p > 0.5).

**Figure 3.**
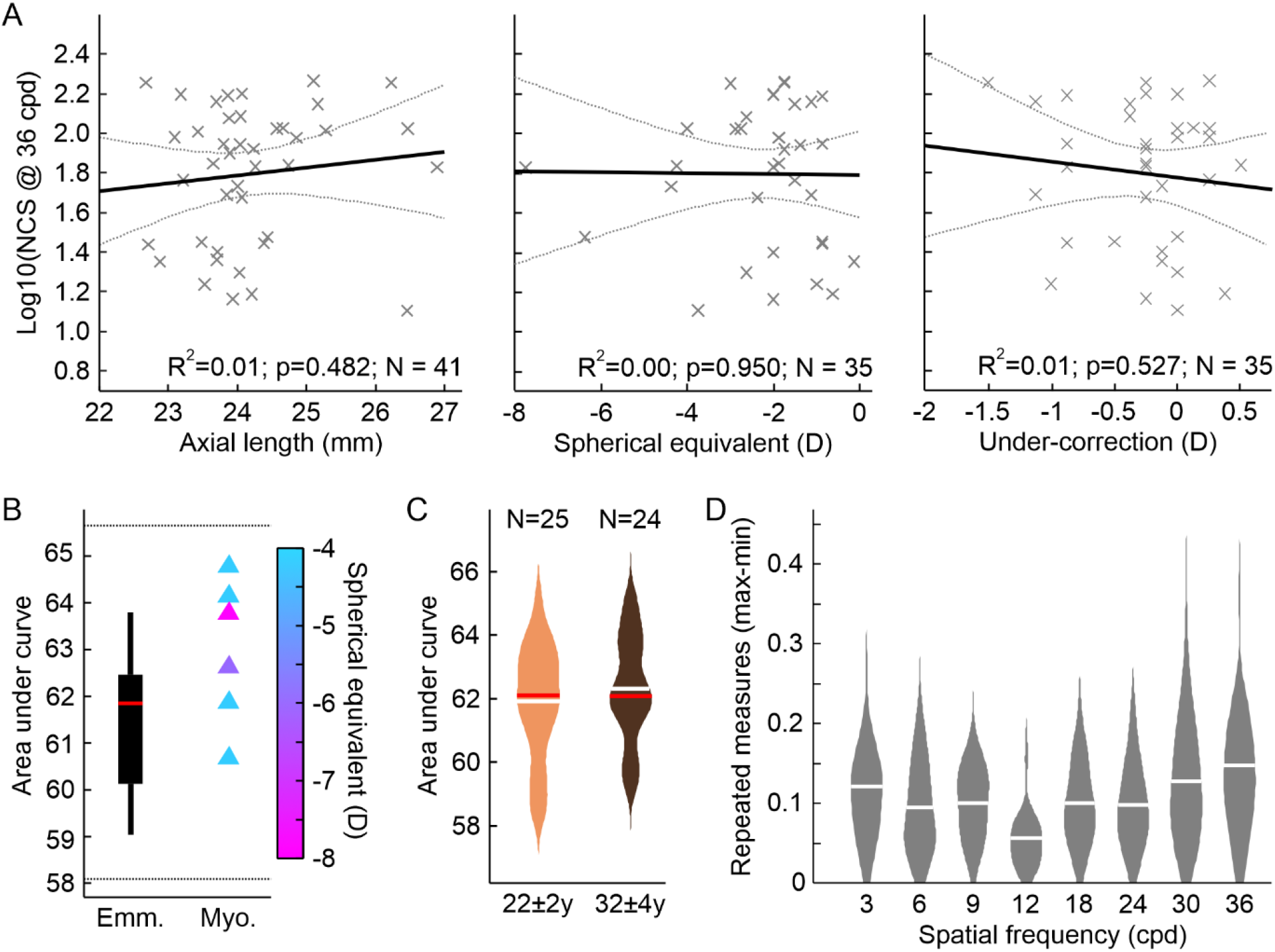
Neural contrast sensitivity is neither correlated with the eye’s refraction nor age. A) None of the tested correlations between observed NCS at 36 cpd and physical parameters of the eye (axial length, refraction given by spherical equivalent, or under-correction) was significant. For axial length NCS measurements were pooled across all three groups, for spherical equivalent and under-correction NCS measurements were pooled from both myopic groups. Grey dotted lines indicate 95% confidence intervals. B) The AUC comparison between emmetropes (SE > -0.5 D; N = 11) and high myopes (SE ≤ -4 D; N = 6) revealed a non-significant difference (p = 0.06), but significant equivalent NCSF (p = 0.03). TOST limits indicated by the dashed lines. C) Division of the pooled data set into two age groups (22±2 years and 32±4 years) showed no significant difference (p=0.50), but significant equivalence (p<0.001). D) 93 % of repeated NCS measurements at a given spatial frequency showed a maximum difference of 0.2 log10 NCS or less between the highest and smallest value recorded. White lines indicate the average, red lines the median value.

Based on this observation, the NCS curves of the highly myopic participants (N = 6), here defined by a refraction ≤ -4 D, from both myopic groups were tested in comparison with the emmetropic group. There was no significant difference between the AUCs of the two groups (p = 0.06), instead both were significantly equivalent (p < 0.05), given the above-defined limits of equivalence (Fig. 3B).

Testing any age-related influence on the NCS within 20 and 40 years, the data set was divided into two subgroups. One group with an average age of 22 ± 2 years (N = 25) and the second (N = 24) with 32 ± 4 years (Fig. 3C). Statistical comparison of the two groups showed no significant difference, but significant equivalence (p = 0.50, and p < 0.001, respectively).

### Low variability for repeated measurements

To assess the repeatability and therefore reliability of the method, the spread of the three repeated NCS measurements per SF was analyzed given by the difference between the maximum and minimum value (Fig. 3D). Overall, 95% of the repeated measurements had a spread of 0.21 log10(NCS). The average spread decreased from 0.12 (STD: ±0.09) at 3 cpd to its lowest value of 0.06 (±0.04) at 12 cpd and increased for higher SFs up to 0.15 (±0.10) at 36 cpd.

### Estimated cut-off frequency, Nyquist limit, and visual acuity

The cut-off frequency based on the fit function for the healthy participants would be approximately 79 cpd, just slightly higher than the foveolar Nyquist limit of two participants (emmetropic: 70.7 cpd, myopic: 67.3 cpd) representatively analyzed (Fig. 4). The FrACT-derived cut-off frequency was lower than the Nyquist limit and with a comparable offset in both analyzed eyes (55.4 cpd and 51.0 cpd, for the emmetropic and the myopic participants, respectively).

**Figure 4.**
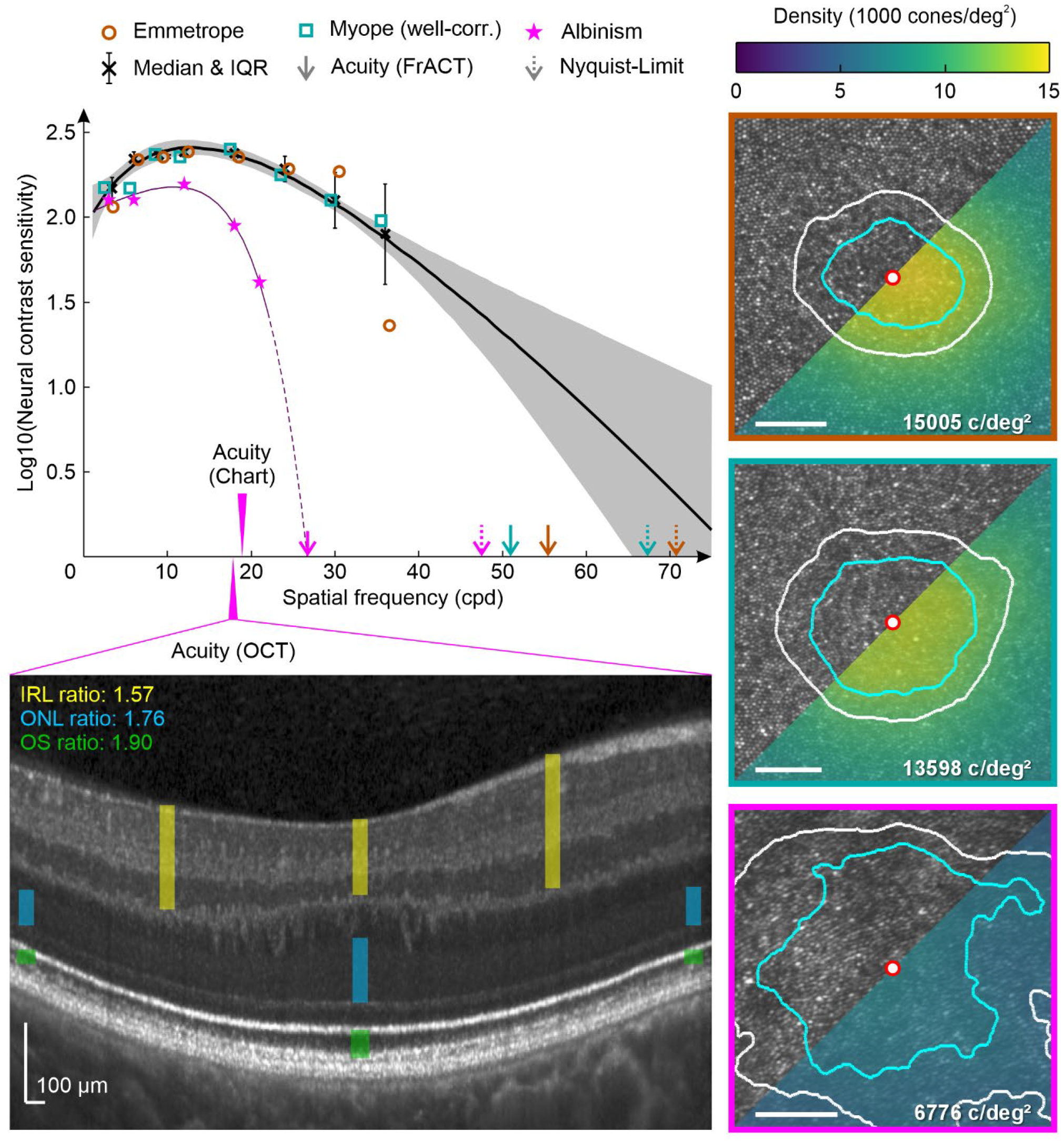
The general healthy neural contrast sensitivity curve and comparison with an albinism patient. At each tested spatial frequency all available NCS measurements were pooled and the resulting Medians (black Xs, errorbars indicate the respective interquartile range) were fitted with a quadratic function (black line), the grey lines show the 95% confidence interval of the fit. Dotted lines indicate the extrapolation of the NCS curve for higher spatial frequencies not tested here. The estimated cut-off frequency (intersection of the fitting line with the abscissa) matches with the Nyquist limit of two study individuals. The Nyquist limit was obtained from foveolar cone densities based on AOSLO imaging (right side). The best-corrected visual acuity measured with FrACT for these two healthy individuals was slightly reduced compared to the Nyquist limit. For the albinism data a rational function of degree (1,2) yielded the best fit. The estimated cut-off frequency was closely related to the best-corrected visual acuity (FrACT), but significantly less than the AOSLO based Nyquist limit. Using the method recently proposed by Woertz et al. to predict the visual acuity from OCT images in albinism patients^34^ matches with the Snellen chart visual acuity but is considerably less then the estimated cut-off frequency and FrACT visual acuity.

### Neural contrast sensitivity for an albinism patient

The normal NCS curve from the pooled healthy data set (see above) was then compared to the NCS recording of an albinism patient with known foveal hypoplasia. Only for the lowest SF tested (3 cpd), the NCS measurements of the albinism patient overlapped with the normal curve. At a SF of 12 cpd, the NCS curve for the albinism patient reached its maximum, like the normal NCS function, but was significantly lower (albinism median NCS = 2.19; normal NCS = 2.41). While the normal NCS shows a rather flat decreasing slope after the peak, the albinism’s NCS decreases steeply, such that the NCS at a SF of 24 cpd could not be tested in our system, limited by the maximum available contrast of 6.3 % (= log10(NCS) of 1.2).

In a final step, the predicted cut-off frequency for the albinism patient, estimated by fitting a Rational-1-2 function (R^2^ = 0.99), was compared to the Nyquist limit based on the individual foveolar cone mosaic (Fig. 4, lower right image). The predicted cut-off frequency from the NCS-fit (26.7 cpd) was much lower than the Nyquist limit of the foveola (47.5 cpd). But it matched exactly with the cut-off frequency obtained by converting the minimum angle of resolution from visual acuity testing. However, when using the minimum angle of resolution from a commonly used letter chart acuity test, this estimated cut-off frequency was lower (18.9 cpd) compared to the albinism NCS-fit prediction. A recently proposed method by Woertz et al. was applied to predict visual acuity based on structural features of the individual retina obtained via optical coherence tomography (OCT).^34^ This OCT-derived cut-off frequency (17.8 cpd) was again lower than the NCS-based estimate, but close to the letter chart acuity.

## Discussion

Motivated by a world-wide increasing prevalence of myopia and the associated increasing numbers of potentially uncorrected and under-corrected myopes, the aim was to test whether neural contrast sensitivity (NCS) is affected and possibly reduced due to long-term neural adaptation to myopic blur. Using an improved interferometric setup that enabled aberration-free stimulus presentation in a larger number of eyes, we found equivalent NCS functions for emmetropes, well-corrected myopes, and under-corrected myopes.

Previous studies using a similar approach or adaptive optics (AO) to correct optical errors, reported degraded NCS in keratoconus patients because of long-term adaptation to optical errors.^38^ Other studies reported orientation specific NCS functions for keratoconus patients^39^ as well as in cases of astigmatism^40^ correlated with the individual asymmetrical optical aberrations of their eye. For myopic participants, a decreased visual resolution after AO correction compared to emmetropic participants was reported.^41,42^ A possible explanation was that such a reduction in sensitivity could result from retinal stretching in myopia leading to a reduced foveolar cone density^43^. Latest advances in retinal imaging made it possible to resolve the foveolar cone mosaic and it was reported that in fact, myopic and thus longer eyes have on average slightly higher angular foveal cone densities compared to shorter, emmetropic eyes.^44,45^ Indeed, the here applied contrast detection task revealed that NCS functions for emmetropes and well-corrected myopes were significantly equivalent. Additionally, a specific comparison of the NCS functions between high-myopes (here defined by a spherical equivalent ≤ -4D) and emmetropes, also showed a significant equivalence. This is in line with the finding that foveal acuity, assessed via an interferometer, was similar in terms of angular units between myopes and emmetropes.^22^ The reason for the different observations from AO and interferometer based studies is most likely that the AO stimulus is affected by unknown residual uncorrected wavefront errors and retinal magnification, while the interferometer stimulus is largely independent of the eye’s aberrations and axial length.^46^

Based on these observations, a reduced NCS could be expected for under-corrected myopes. At a far viewing point, an under-correction of 1 D or more largely diminishes the modulation transfer function, leading to a significant signal reduction of medium spatial frequencies (SFs,10 to 20 cpd) and complete elimination of high SF. However, this was not the case and a significant equivalence of the NCS functions between emmetropes and under-corrected myopes was found. The observation of a result that contradicts the hypothesis could be explained as follows: While the optical aberrations of keratoconus patients can never be fully corrected in daily life, myopes have their distance point close by, allowing them to see clearly at short distances even without correction. Thus, during near work, the high spatial frequency channels are sufficiently stimulated and remain active.

Across all participants, the NCS peak was found at about 12 cpd (Fig. 2A and B), which is higher compared to previous studies reporting peak NCS at 8 cpd or 10 cpd.^17,18^ A small subset of emmetropic participants was tested at a SF of 9 cpd, yielding equal NCS measurements for 9 and 12 cpd. This supports, even though not specifically measured, the peak SF to be rather in between the two SFs. It can be furthermore estimated that the 12 cpd measurements are closer to the true peak sensitivity than the 6 cpd from the observation that for 12 cpd the standard deviation across the tested participants (Fig. 2D), as well as the spread for repeated measurements in the same participants (Fig. 3D), was lowest.

The absolute NCS values for low SFs (3 and 6 cpd) were similar to those reported by Campbell and Green.^17^ In contrast, the observed NCS at high SFs (≥ 18 cpd) was higher than the previous reports, but with a similar slope for increasing SFs. Compared to Williams,^23^ the currently reported NCS values are slightly higher than the values reported for the most sensitive participants. A possible explanation for this could be that the center wavelength in our setup was 550 nm compared to 630 nm in the former studies. 550 nm light stimulates L- and M-cones equally, while 630 nm activates L-cones more than M-cones. It was shown that monochromatic NCS measurements at 543 nm are increased by about 0.15 log10 NCS compared to 623 nm,^47^ which corresponds to the offset between measured and reported (Williams) NCS for SFs from 12 to 24 cpd.

The observation of similar NCS functions measured for such a large cohort, contradicts earlier reports calculating the NCS function, stating it to be highly variable and individual, especially at medium SFs between 3 and 20 cpd.^48^ For high SF (≥ 30 cpd) a striking variability increase between individual NCS values was observed, with the interquartile range increasing from about 0.1 (≤ 30 cpd) to 0.6 at 36 cpd. Such idiosyncratic fluctuation of NCS at high SFs has been observed before with a difference of 0.4 log10 units between the highest and lowest sensitivity values.^17,23^ The refractive state of the eye, axial length, and under-correction showed no correlation with NCS measurements at 36 cpd, suggesting these factors are not the source of the observed effect. Another possible explanation is a masking effect due to vitreous opacities or an instable tear film. For example, it was reported that a stable tear film increases visual acuity, while acuity decreases for dry eyes.^49^ Although an interferometric system is supposed to bypass the eye’s optics, it is unclear to what extent the laser beams interfere with particles in the tear film, possibly degrading stimulus quality to an unknown extent.

Hypothetically, the NCS measurements especially for high SFs could have been affected by adaptation, due to the unusual aberration-free stimulus projection, resulting in an underestimation of the true NCS.^50,51^ For example, a 15 sec adaptation time to a grating stimulus being 0.75 log10 units above threshold led to a threshold increase of contrast sensitivity by 0.2 log10 units.^52^ With an average number of 32 trials tested here, the total summed up stimulus duration would be 16 sec. However, the individual stimulus presentation duration of 500 msec was extremely short compared to this. Additionally, the stimulus presentation was followed by a period of at least 1 to 2 sec without a stimulus, depending on the response time. Furthermore, because of the implementation of QUEST to determine the threshold, most of the stimuli were displayed with a contrast close to the threshold, additionally minimizing any influence of adaptation. Given these procedures, any visual adaptation to the stimulus which could have led to an underestimation of the NCS can be neglected.

For a resolution task it was recently shown that with adaptive optics correction, visual acuity is correlated to the retina’s Nyquist limit.^54^ However, the estimated cut-off frequency from a rational fit exceeds the Nyquist limit of a healthy retina (see Fig. 4). For a detection task, which was used here, this observation has been reported earlier^18,53^ and could by explained by aliasing, which allows stimulus discrimination even at SFs beyond the Nyquist limit.^18,55,56^ However, for central vision contrast thresholds between a resolution and detection task are similar up to a SF of 50 cpd^57,58^, meaning that the here measured NCS for SPs between 3 and 36 cpd are not affected by aliasing.

Under these circumstances, it is surprising that for the albinism patient, a steep drop in NCS beyond 12 cpd was observed. In this patient, the estimated cut-off frequency was much lower than the retina’s Nyquist limit (about 27 cpd compared to 47 cpd). A related observation was reported earlier for albinism patients, with visual acuity not reaching the Nyquist limit.^59^ A possible explanation for this is that in the foveal region of a healthy retina, each cone connects with a single midget retinal ganglion cell, while in albinism the absence of a true fovea goes along with a lack of a preserved private line pathway.^60^ However, the estimated cut-off frequency matched the visual acuity determined via FrACT closely, while these two read-outs were higher than the letter chart and an OCT-based visual acuity estimation. Due to the underlying algorithm of FrACT, placing the stimulus always at the estimated threshold, it was reported earlier that FrACT visus values are higher on average compared to the traditionally used letter chart.^61,62^ Considering the here reported findings for NCS, the FrACT values are more closely related to the true resolving capability of the individual retina, especially in albinism.

When the pooled data set was divided into two subgroups (22±2 and 32±4 years old), to test age-related influences, the NCS functions of the two groups were significantly equivalent. This is contradictory to the previous finding that the NCS deteriorates with each decade,^63^ but confirms another study, where only a small difference in NCS functions between younger and elderly participants (average age 21 and 68 years) were observed.^64^ Thus, further research is needed to specifically investigate NCS in the healthy aging eye.

In conclusion, an interferometric system with an improved optical design by utilizing a spatial light modulator enabled the measurement of neural contrast sensitivity in 49 untrained participants. Robust neural contrast sensitivity functions were observed, unaffected by myopia and under-correction. Thus, long-term under-corrected myopes will be able to achieve normal visual acuity as soon as their refractive errors are sufficiently corrected. Because of the changed optical design and shorter wavelength compared to classic interferometer-based NCS measurements, the currently reported absolute NCS values are slightly higher, but in accordance with previous findings. Therefore, a generally applicable NCS function for young healthy eyes is proposed. For two healthy participants the estimated cut-off frequency is close to the Nyquist limit and higher than the best corrected visual acuity, demonstrating that human central vision in the healthy eye is mainly limited by the eye’s optics. A first comparison of the normal NCS function with NCS values obtained from an albinism patient revealed a significant NCS degradation. With the estimated cut-off frequency being significantly lower than the Nyquist limit, this leads to the conclusion that vision in albinism is mainly limited by neurological factors.

## Acknowledgements

The authors thank all participants for their contribution in this study.

## Author contributions

Conceptualization: N.D., K.B., and J.M.; Investigation: L.R., J.M., N.D., and J.A.; Writing – Original Draft: N.D.; Writing – Review & Editing: All authors; Visualization: N.D.; Funding Acquisition: S.W. and W.M.H.; Resources: S.W. and W.M.H.; Supervision: N.D., K.B., W.M.H., and S.W.

## Ethics approval

The study adhered to the tenets of the Declaration of HELSINKI (2013). The ethics authorization to perform the measurements was granted by the Medicine Faculty Human Research Ethics Committee from the University of Tübingen with the ID 616/2022BO2 and 094/2024BO1. Prior to data collection, the experiment was explained in detail to the participants, and written informed consent was collected from each participant. All data were pseudonymized and stored in full compliance with the principles of the Data Protection Act GDPR 2016/679 of the European Union.

## Lead contact

Further information and requests for resources should be directed to and will be fulfilled by the lead contact, Siegfried Wahl.

